# Foot perfusion. Insights from an anatomically detailed arterial network model

**DOI:** 10.64898/2025.12.05.692561

**Authors:** Malene Bisgaard, Caterina Dalmaso, Jens Vinge Nygaard, Helle Precht, Kim Christian Houlind, Lucas Omar Müller, Pablo Javier Blanco

## Abstract

Peripheral artery disease currently affects over 202 million people worldwide. The ankle-brachial index is one of the most used measurements to assess a reduction in blood flow to the foot, but it is not able to characterise tissue perfusion. Information on tissue perfusion can be obtained, among others, through an MRI scan, which is time-consuming and can be painful if induction of ischemia is warranted for the scan. As an alternative, we model foot perfusion during a cuff-induced ischaemia test to characterise how occlusions in foot arteries affect perfusion in foot regions. Simulations are not patient-specific at this stage, and are conducted on a 1D arterial network model which includes 154 foot and calf arterial segments, providing a realistic description of the topology of the foot arterial vasculature. A baseline model characterizes perfusion in angiosomes under healthy conditions, which is then modified to reflect 42 pathological scenarios by introducing occlusions and different levels of collateral impairment. Results show a marked influence of collateral impairment on angiosome perfusion under the condition of a single-artery occlusion, highlighting the role of blood redistribution. If two feeding arteries are occluded, perfusion markedly decreases at all collateral impairment levels due to the severe reduction in incoming blood flow. These results provide a bridge between the angiosome-targeted and “best-vessel” strategies for revascularization, showing that both can be correct depending on collateral sufficiency.

**Graphical Abstract:** 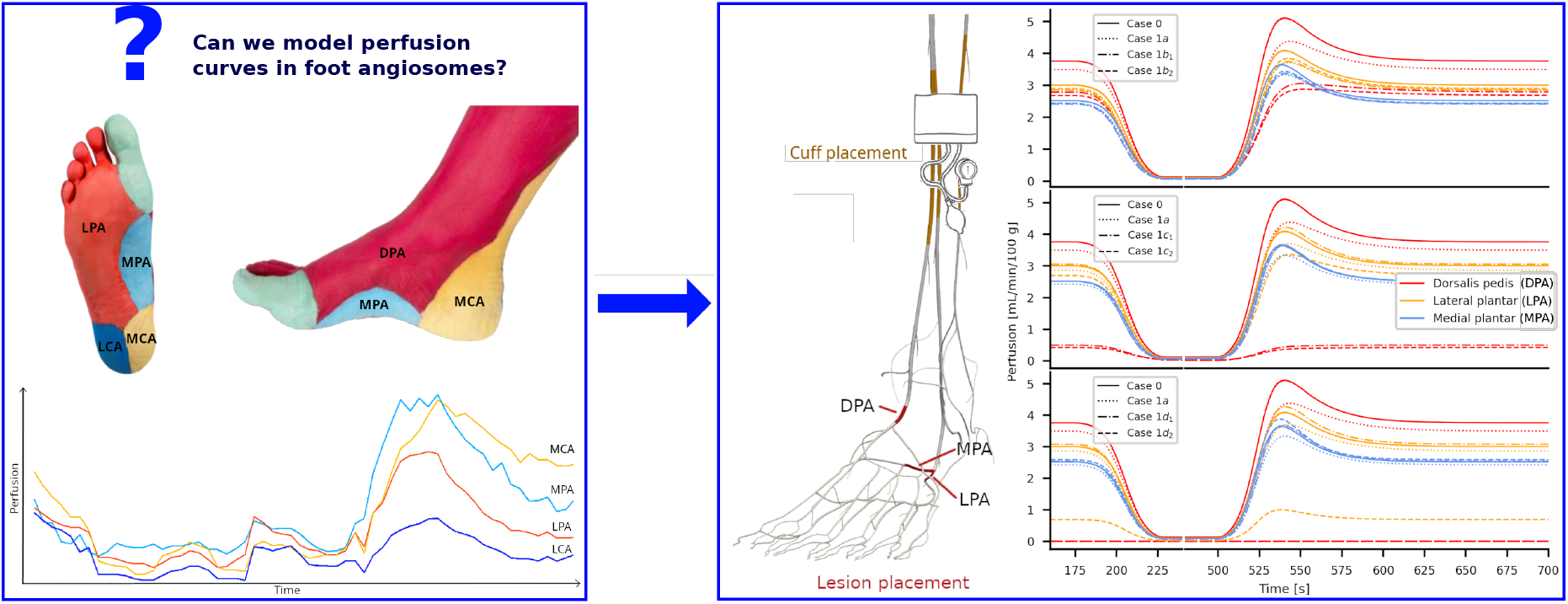

**Highlights:**

- We propose a computational model of foot perfusion during a cuff-induced ischaemia test that allows the assessment of perfusion in each angiosome of the foot
- We study how occlusions in the main feeding arteries of the foot impact tissue perfusion
- We highlight the role of collateralization if adequate inflow is maintained
- Results show that both angiosome-targeted and “best-vessel” strategies for revascularization can be correct depending on arch patency and collateral sufficiency

## 1. Introduction

More than 202 million people worldwide suffer from peripheral artery disease (PAD) (Song et al., 2019). For some patients with PAD, the disease manifests as critical limb-threatening ischemia (CLTI): their mortality risk increases (Van den Berg, 2018), and it is necessary to restore perfusion to damaged tissues to avoid amputation. To guide vascular surgeons in selecting the best treatment, patients will have an ankle-brachial index (ABI) or a toe-brachial index (TBI) measured. In addition, the leg vessels will be imaged using ultrasound, subtraction angiography, computed tomography, or magnetic resonance imaging (MRI). These examinations allow surgeons to visualize occlusions and stenoses, as well as the largest collateral vessels, thereby providing information on blood flow to the leg and foot. Based on them, the best revascularization or conservative treatment is decided (Norgren et al., 2007). Studies suggest that it is important to consider the angiosome theory to ensure revascularisation of the damaged tissue when treating patients with CLTI (Špillerová et al., 2017; Neville et al., 2009). For patients with ulceration, locating the foot area that needs reperfusion is possible. However, if a wound covers more than one angiosome, the perfusion status in each angiosome would be relevant. Furthermore, perfusion measurement could be relevant to evaluate treatment effect. In the last few years, even though it has been possible to image the perfusion in all modalities that produce angiography (Boonen and Aerden, 2023; Caroca et al., 2021; Reekers et al., 2016; Souza et al., 2024), these measurements have not yet become a clinical standard. Several reasons are behind this: new software is required for generating and evaluating the images, staff need to be trained, and examinations can be time-consuming and painful for the patient.

As an alternative, computer simulations may provide estimates about the perfusion level in each foot angiosome based on information from the angiography. This study aims to characterize how occlusions in the main feeding arteries impact perfusion, in order to explore the applicability of the angiosome-targeted and “best-vessel” paradigms for revascularization at different degrees of collateral impairment. To this end, it is necessary to employ models endowed with a sufficiently detailed description of the vascular topology of the leg and the foot, for them to capture both direct and alternative feeding routes to the different angiosomes (Varela et al., 2017). In particular, in addition to the main feeding arteries, such a model should include arterial-arterial connections such as the plantar arch and the distal peroneal branches. As a consequence, we chose to employ the Anatomically-Detailed Arterial Network (ADAN) model (Blanco et al., 2014b,a), which includes 2142 vessels, comprising almost every named arterial vessel. In particular, the topology of the vascular architecture in the lower limb is extremely realistic, with 154 vascular segments in the foot and calf.

The rest of the work is structured as follows. In Section 2, we describe the clinical question and the mathematical model employed for simulations. Section 3 collects simulation results for the baseline and pathological scenarios, which are discussed in Section 4. Finally, in Section 5 we summarize our main findings.

## 2. Methods

In this section, we provide an overview of the clinical question being the basis of this study, together with explanations of the mathematical model employed for simulations.

### 2.1. Clinical problem

Here, we outline the main features of the clinical problem, describe the expected behaviour of foot perfusion during cuff-induced ischaemia tests, and illustrate the two main theories guiding the choice of vessels that need revascularization.

#### 2.1.1. Foot angiosomes

Angiosomes can be defined as areas of tissue (skin, subcutaneous tissue, fascia, muscle and bone) supplied and drained by specific arteries and veins. According to the most common subdivision, the foot can be decomposed into 5 different angiosomes, perfused by the medial plantar artery (MPA), lateral plantar artery (LPA), dorsal pedal artery (DPA), medial calcaneal artery (MCA), and lateral calcaneal artery (LCA) (Houlind and Christensen, 2013). Furthermore, the first toe is supplied by DPA, MPA, and LPA. In Figure 1 these angiosomes are depicted, respectively, in light blue, orange, red, yellow and blue. The presence of a well-developed collateral network in the foot provides alternative perfusion/drainage routes, allowing angiosomes to receive blood from neighbouring regions in the case of obstructive disease.

**Figure 1.**
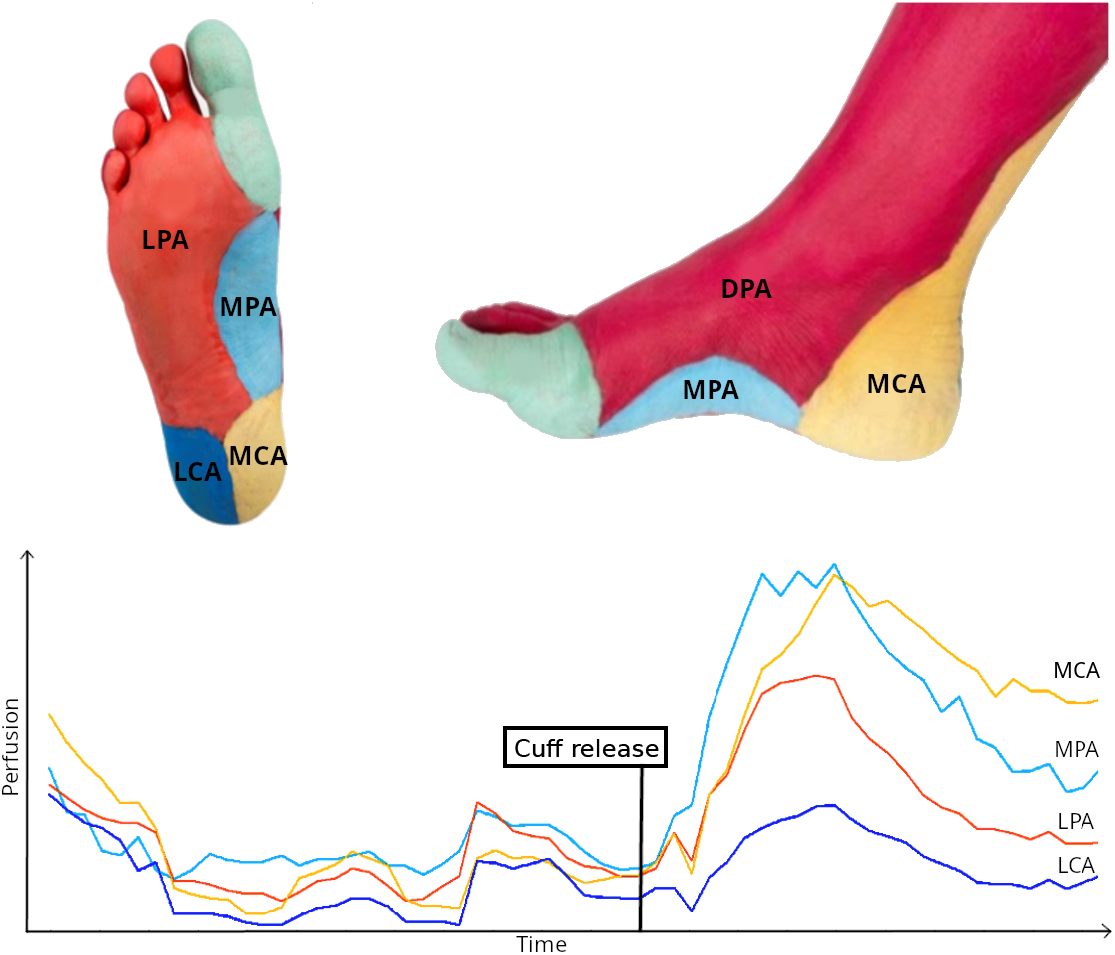
Foot angiosomes and example of perfusion curves obtained using a Flow-sensitive Alternating Inversion Recovery (FAIR) sequence. Light blue: angiosome perfused by the MPA; orange: angiosome perfused by the LPA; red: angiosome perfused by the DPA; yellow: angiosome perfused by the MCA; blue: angiosome perfused by the LCA. Adapted from (Houlind and Christensen, 2013).

#### 2.1.2. Revascularization guidelines and clinical practice

In the case of CLTI patients, perfusion can be restored to damaged tissues via revascularization. Two main theories have emerged regarding which vessels should be treated in order to provide the best outcome. Some clinicians claim that perfusion to the ischemic angiosome should be obtained by direct revascularization of the relevant feeding artery (Houlind and Christensen, 2013). An alternative view claims that one should aim for revascularization of the best preserved vessel that crosses the ankle, regardless of whether or not this vessel provides direct flow to the relevant angiosome (Norgren et al., 2007).

#### 2.1.3. Perfusion in a cuff-induced ischemia test

In recent years, various imaging modalities have demonstrated potential for assessing perfusion or oxygenation in the feet. These modalities include positron emission tomography–computed tomography (PET-CT) (Christensen et al., 2024; Burchert et al., 1997), digital subtraction angiography (Reekers et al., 2016), computed tomography (Boonen et al., 2023), ultrasound (Sommerset et al., 2019; Souza et al., 2024), near-infrared spectroscopy (Meertens et al., 2021), indocyanine green fluorescence imaging (Fang et al., 2022; Tange et al., 2023, 2024), thermal imaging (Gatt et al., 2018; Petrova et al., 2018), and MRI (Caroca et al., 2021). However, to the best of our knowledge, none of these techniques have been implemented in routine clinical practice. A promising method for measuring foot perfusion is arterial spin labeling (ASL), measured with MRI. This technique enables quantitative perfusion measurements without the use of gadolinium-based contrast agents (Leithner et al., 2008). Since the ASL signal is proportional to perfusion, which is low at rest in the foot, this approach requires a cuff-induced ischaemia test, to increase perfusion for a short time period (Lopez et al., 2015). In particular, ASL allows the generation of perfusion curves, as illustrated in Figure 1, depicting baseline perfusion levels and time to peak (TTP), defined as the interval between cuff release and the attainment of maximum perfusion.

### 2.2. Mathematical model

In the following subsections, we outline the modelling setup adopted for simulations. In particular, we illustrate the main features of the arterial network model, and describe the myogenic local autoregulation model, incorporated in order to reproduce the transient hyperaemic response following cuff deactivation (reactive hyperaemia). In particular, an impaired reactive hyperaemia indicates microvascular dysfunction (Rosenberry and Nelson, 2020), and is related to the inability of the vessels to display an autoregulatory response.

#### 2.2.1. Arterial network model

We adopt the Anatomically-Detailed Arterial Network (ADAN) model developed by Blanco et al. (2014a,b), which provides a comprehensive source of anatomical detail regarding the topology of the vascular architecture in the lower limb, notably the foot. In particular, the vascular network includes 154 foot and calf arterial segments (each segment being defined by the presence of a bifurcation). This anatomical detail provides a baseline (healthy/lesion-free) characterization of blood flow to the different vascular territories of the foot (see Figure 2). All the parameters for the ADAN model are calibrated as reported in Blanco et al. (2014a, 2020); Müller et al. (2023). The complete model dataset is publicly available at http://hemolab.lncc.br/adan-web. The baseline model is then modified to reflect 42 pathological scenarios by introducing occlusions in different arteries of the foot.

**Figure 2.**
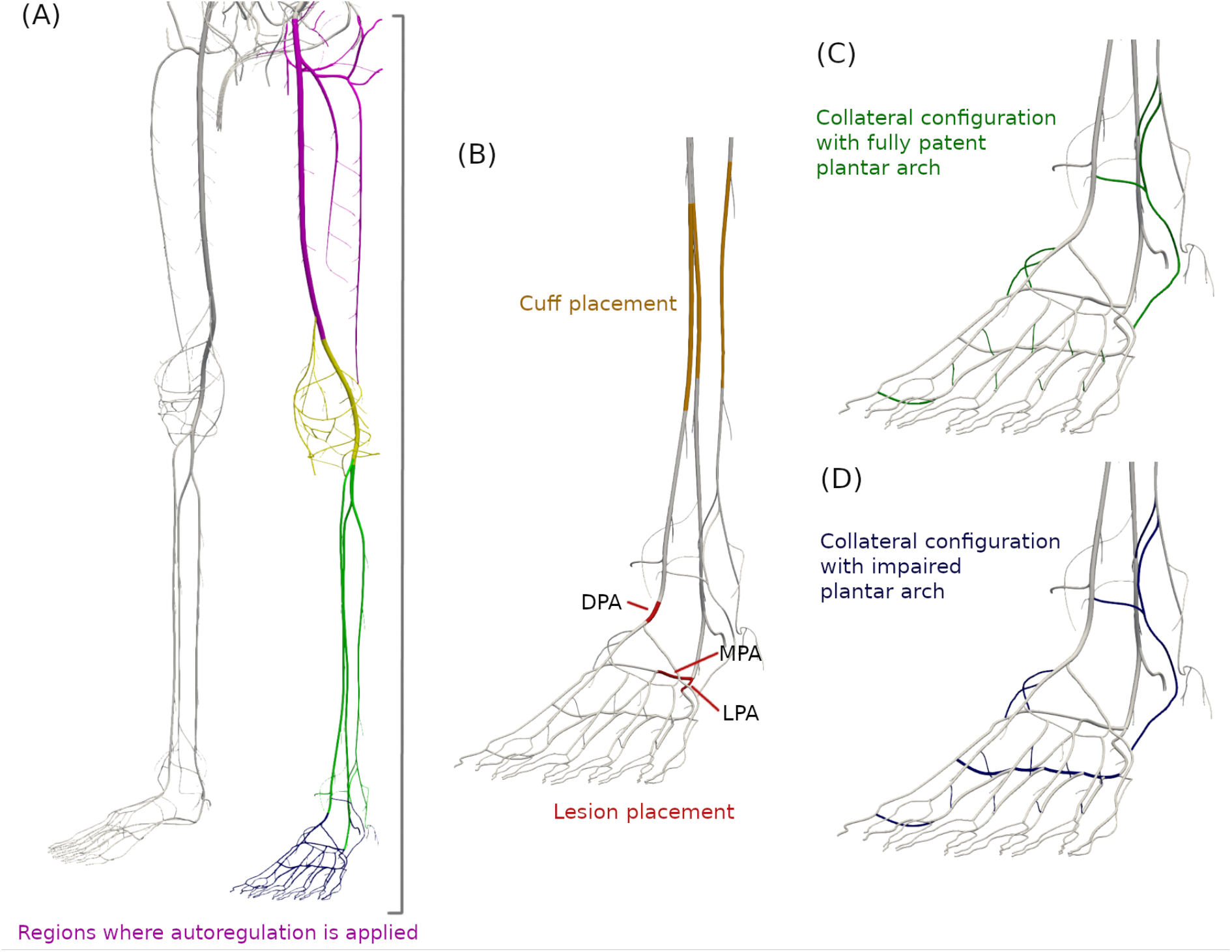
Arterial vasculature in the leg. Panel (A): regions where the autoregulation model is applied (each color corresponds to a different parametrization, see Table 2); Panel (B): cuff and lesion placement (DPA, MPA, LPA stand, respectively, for dorsalis pedis, medial plantar, lateral plantar artery); Panels (C) and (D): foot collaterals.

Blood flow simulations are based on the one-dimensional (1D) theory of the incompressible flow of a fluid in compliant vessels. The governing equations describe the pressure *P*, flow rate *Q*, and lumen area *A* as a function of time *t* and the longitudinal coordinate *x* (Hughes and Lubliner, 1973), and read as follows

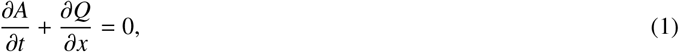

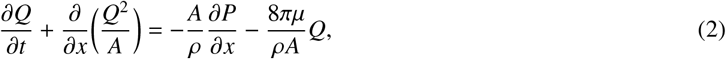

where *ρ* = 1.04*g/cm*^3^ is the blood density, and *µ* is the fluid viscosity. Blood viscosity is set to *µ* = 0.04*P* everywhere except for perforator arteries, where *µ* = 0.01*P*. The vessel wall is considered to be a non-linear viscoelastic material whose governing equation is

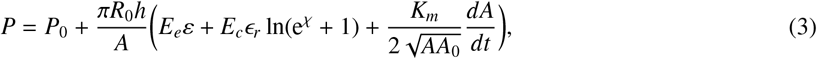

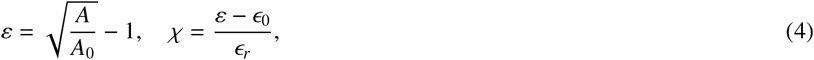

where *R*_0_ and *A*_0_ are the reference lumen radius and area at the reference pressure *P*_0_, *h* is the arterial wall thickness, *ϵ*_0_, and *ϵ*_*r*_ describe the mean and standard deviation of the fibre recruitment distribution, *E*_*e*_ and *E*_*c*_ stand for the effective elastic moduli of the elastin and collagen fibres, respectively, and *K*_*m*_ is the viscoelastic coefficient. Material parameters of the tube law (*E*_*e*_, *E*_*c*_, *K*_*m*_), the parameters describing collagen fibres (*ϵ*_0_, *ϵ*_*r*_), and wall thickness *h* are computed as in Blanco et al. (2014a); Müller et al. (2023). The pressure of the reference state is *P*_0_ = 10^5^*dyn/cm*^2^.

The principles of mass and total pressure conservation determine coupling conditions at vascular junctions in the network (Müller et al., 2016). For *N* converging vessels at a junction, this implies

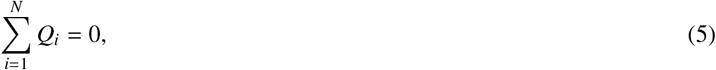

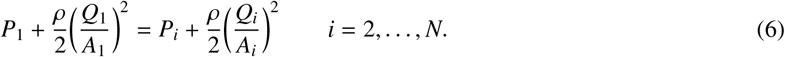

The total pressure conservation equation (6) is modified to simulate the inflation and deflation of a cuff, which is modeled as a two-vessel junction where total pressure is not conserved. In particular, setting vessel 1 as the vessel sharing its outlet with the junction, we enforce that

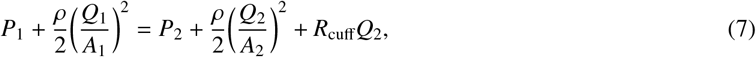

Here, *R*_cuff_ changes depending on the state of the cuff: it is null while the cuff is deactivated, maximal while the cuff is activated, and varies according to an exponential profile during cuff activation and deactivation

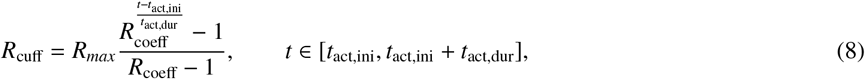

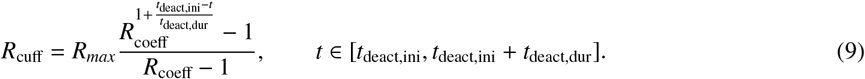

Here, *t*_*act,ini*_ and *t*_deact,ini_ are the times at which cuff activation and deactivation begin, and *t*_act,dur_ and *t*_deact,dur_ are the durations of the activation and deactivation phases. *R*_coeff_ is a coefficient that determines the speed of activation/deactivation, and *R*_*max*_ is the maximum resistance reached when the cuff is fully activated.

Peripheral boundary conditions are modelled using 3-element Windkessel elements, that is

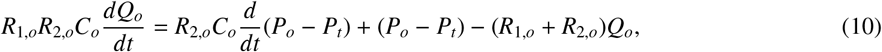

where *P*_*T*_ is the reference terminal pressure, *R*_1,*o*_, *R*_2,*o*_, and *C*_*o*_ are the parameters that represent peripheral resistances and compliance downstream of each terminal vessel. The resistance of a peripheral bed is *R*_*tot,o*_ = *R*_1,*o*_ + *R*_2,*o*_, with *R*_1,*o*_ = 0.2*R*_*tot,o*_, *R*_2,*o*_ = 0.8*R*_*tot,o*_. Total peripheral compliance is 10% of arterial tree compliance, and is distributed among terminals according to the blood flow fraction through them (Blanco et al., 2014a). These 0D models reproduce the effects of the remaining vasculature (microcirculation and venous circulation).

Occlusions are modelled by enforcing no-flow conditions across them.

#### 2.2.2. Autoregulation model

We implement a local myogenic flow regulation model based on Ursino and Giannessi (2010), which provides feedback when the flow rate departs from a homeostatic state, defined here by the baseline model state mentioned above. In particular, this control model acts upon peripheral resistances and compliances whenever the flow through the corresponding peripheral Windkessel models is affected as a consequence of modifications in the vascular network. This allows us to reproduce the hyperaemic response after cuff opening, caused by vasodilation, as well as the ability of the vessels to preserve an adequate flow rate if their calibre is reduced due to pathological conditions. Without such a model, blood flow would be restored to its baseline value immediately after cuff release.

The effect *e*_*o*_ of autoregulation at the level of each terminal *o* in region *reg* is governed by

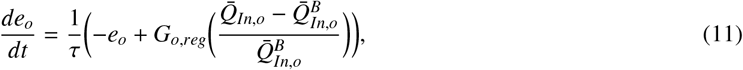

where *τ* is the time constant of this low pass dynamic, *G*_*o,reg*_ is its static gain, and 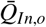 and 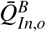 are, respectively, the cardiac cycle-averaged flow rate and baseline flow rate at the midpoint of the 1D vessel coupled to the 0D terminal. The static gain *G*_*o,reg*_ at each terminal *o* in region *reg* is computed from the total gain *G*_*reg*_ of autoregulation, which is considered proportional to the flow in each region. Once the control action *e*_*o*_ is available, it is used to modify the terminal compliance *C*_*o*_ (see equation (10)) through a sigmoidal relationship as

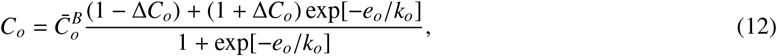

with 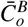 the baseline terminal compliance,

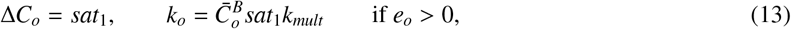

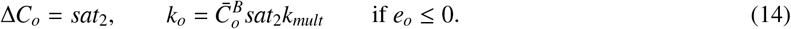

Here, *sat*_1_, *sat*_2_ are constant parameters that define the upper and lower saturation levels of the sigmoidal curve, and *k*_*mult*_ is a constant parameter that regulates the steepness of the sigmoid. If autoregulation is not active, 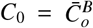. Parameter values, which were determined in order to reproduce the magnitude and time scales of hypereaemic responses observed in the clinical practice, are available in Table 1.

**Table 1:** Parameters employed for the autoregulation model. See color-coded regions in Figure 2.

| Parameter | Regions |  |  |  |
| --- | --- | --- | --- | --- |
|  | 1: pink | 2: yellow | 3: green | 4: blue |
| $\tau$ | 40s | 40s | 40s | 40s |
| $sat_1$ | 0.4 | 0.4 | 0.4 | 0.4 |
| $sat_2$ | 4.5 | 4.5 | 4.5 | 4.5 |
| $k_{mult}$ | 8. | 8. | 8. | 8. |
| $G_{reg}$ | 0.102 | 0.017 | 0.014 | 0.009 |

Equations (11) and (12) imply that a decrease in blood flow below its baseline value 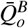, such as the one observed during cuff activation, causes vasodilation, which is modelled through an increase in terminal compliance. Variations in peripheral resistances *R*_2,*o*_ (see equation (10)) are, instead, modelled as

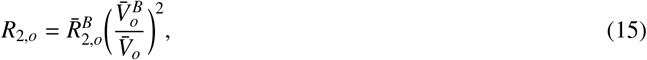

with 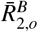 the baseline terminal resistance, and 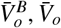 cardiac cycle-averaged baseline and current terminal volumes, respectively. While the cuff is active, terminal resistances temporarily increase due to a decreased blood volume. Right after cuff deactivation, instead, the increase in arteriolar blood volume caused by increased compliances results in a reduction of resistances below their baseline value.

#### 2.2.3. Perfusion estimation

The proposed model allows us to compute the time evolution of flow rates at the level of different foot angiosomes. We can derive reasonable perfusion values in each angiosome from model-generated flow rates by multiplying the flow rate by angiosome weight, which is computed from angiosome volume under the assumption of a constant tissue density of 1*g/cm*^3^. We report in Table 2 the volumes and weights of foot territories in the ADAN model (Blanco et al., 2014b,a).

**Table 2:**
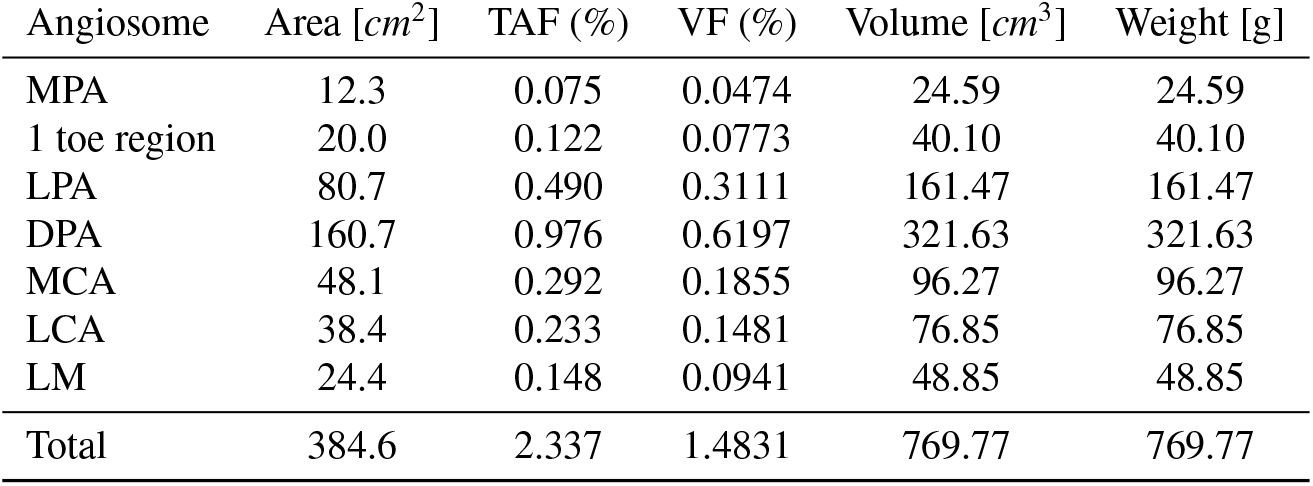
TAF: total area fraction, computed as the ratio between angiosome area and total body surface area (16460*cm*^2^ for the ADAN model); VF: volume fraction, computed as the product between TAF and a scaling factor equal to 0.6347 (Blanco et al., 2014a); MPA: medial plantar artery; LPA: lateral plantar artery; DPA: dorsalis pedis artery; MCA: medial calcaneal artery; LCA: lateral calcaneal artery; LM: lateral malleolar artery.

| Angiosome | Area [$cm^2$] | TAF (%) | VF (%) | Volume [$cm^3$] | Weight [g] |
| --- | --- | --- | --- | --- | --- |
| MPA | 12.3 | 0.075 | 0.0474 | 24.59 | 24.59 |
| 1 toe region | 20.0 | 0.122 | 0.0773 | 40.10 | 40.10 |
| LPA | 80.7 | 0.490 | 0.3111 | 161.47 | 161.47 |
| DPA | 160.7 | 0.976 | 0.6197 | 321.63 | 321.63 |
| MCA | 48.1 | 0.292 | 0.1855 | 96.27 | 96.27 |
| LCA | 38.4 | 0.233 | 0.1481 | 76.85 | 76.85 |
| LM | 24.4 | 0.148 | 0.0941 | 48.85 | 48.85 |
| Total | 384.6 | 2.337 | 1.4831 | 769.77 | 769.77 |

### 2.3. Simulations setup

Simulations were performed to reproduce a cuff-induced ischaemia test, in both a baseline lesion-free scenario, and in multiple pathological cases, defined by occlusions placed in different foot arteries.

### 2.3.1. Cuff protocol

The cuff was applied in all cases to the medial portions of the anterior tibial, posterior tibial, and fibular arteries (see Figure 2, panel (B)). We assume that *R*_*max*_ in equations (8) and (9) is equal to 1*e*^7^[*dyn* ·*s/cm*^5^], so that flow to the angiosomes of the foot during cuff activation is nearly zero. Moreover, we set *R*_coeff_ to 1*e*^4^ and *t*_act,dur_ = *t*_deact,dur_ to 50*s* to obtain reasonably slow cuff activation/deactivation phases. Since after 150*s* all baseline simulations reach a steady-state condition in terms of cardiac-cycle averaged flows, we assume that cuff inflation starts at *t* = 175*s*. The cuff remains fully closed for 275*s*, and, after that, cuff deactivation begins at *t* = 500*s*.

#### 2.3.2. Lesion locations and collateral impairment

Simulations were designed to assess the extent to which the best vessel and angiosome theories are valid at different degrees of collateral impairment, when lesions are present in the main foot arteries. As a consequence, we considered six different lesion configurations, characterized by the presence of at most two occlusions at the level of DPA, MPA and LPA. We performed simulations assuming that foot collaterals are fully patent (scenarios 1*a* - 6*a*) and compared them with simulations with reduced collateral diameters. We distinguished two different collateral configurations, one in which the plantar arch is treated as a collateral vessel (see Figure 2, panel (D)), and one in which it is assumed to be fully patent (see Figure 2, panel (C)). In particular, cases 1*b*_1_ − 6*b*_1_, 1*c*_1_ − 6*c*_1_, 1*d*_1_ − 6*d*_1_ denote scenarios where the plantar arch is assumed to be fully patent, while other collaterals have a diameter that is, respectively, 60%, 30%, and 5% of the original diameter. Cases 1*b*_2_ − 6*b*_2_, 1*c*_2_ − 6*c*_2_, 1*d*_2_ − 6*d*_2_ denote scenarios where all collaterals (including the plantar arch) have a diameter that is, respectively, 60%, 30%, and 5% of the original diameter. The resulting simulation scenarios are summarized in Table 3.

**Table 3:**
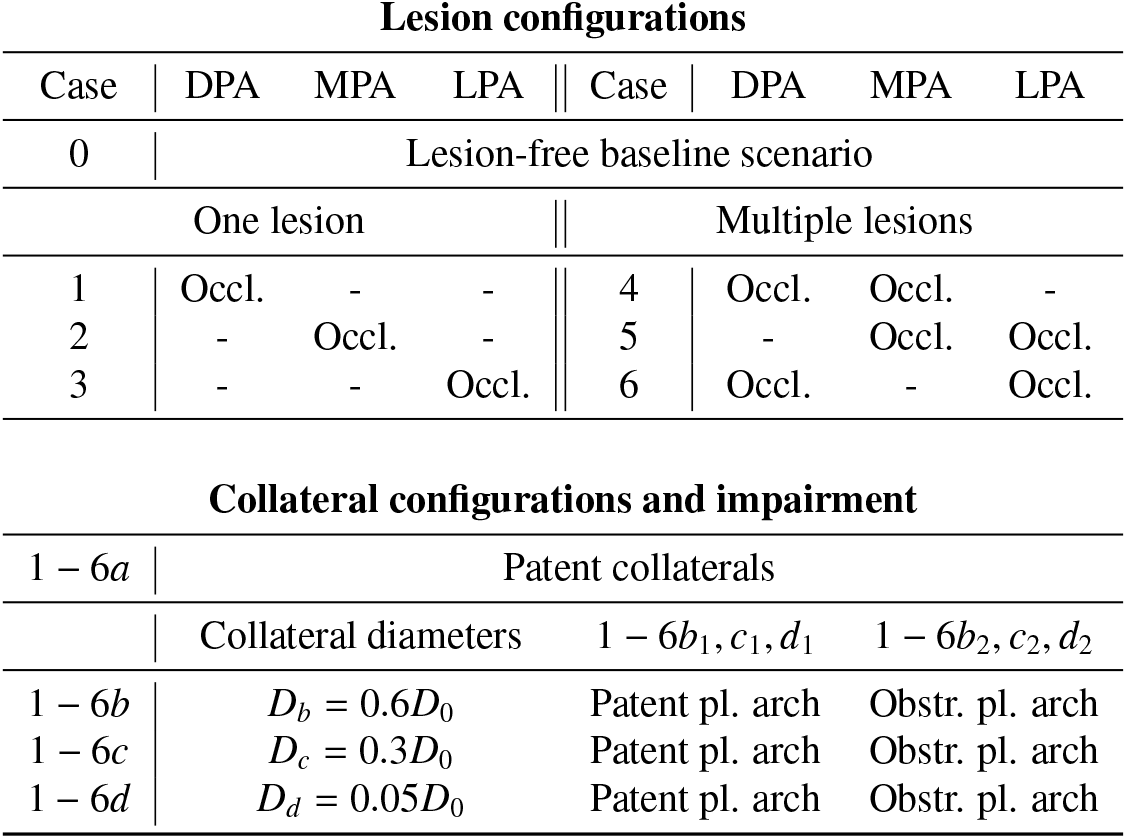
Location of lesions in the dorsalis pedis (DPA), medial plantar (MPA) and lateral plantar (LPA) arteries and collateral configurations (see Figure 2). *D*_0_ is vessel diameter in the baseline lesion-free configuration.

| Lesion configurations |  |  |  |  |  |  |  |
| --- | --- | --- | --- | --- | --- | --- | --- |
| Case | DPA | MPA | LPA | Case | DPA | MPA | LPA |
| 0 | Lesion-free baseline scenario |  |  |  |  |  |  |
| One lesion |  |  |  | Multiple lesions |  |  |  |
| 1 | Occl. | - | - | 4 | Occl. | Occl. | - |
| 2 | - | Occl. | - | 5 | - | Occl. | Occl. |
| 3 | - | - | Occl. | 6 | Occl. | - | Occl. |

| Collateral configurations and impairment |  |  |  |
| --- | --- | --- | --- |
| 1 – 6a | Patent collaterals |  |  |
|  | Collateral diameters | 1 – 6b<sub>1</sub>, c<sub>1</sub>, d<sub>1</sub> | 1 – 6b<sub>2</sub>, c<sub>2</sub>, d<sub>2</sub> |
| 1 – 6b | $D_b = 0.6D_0$ | Patent pl. arch | Obstr. pl. arch |
| 1 – 6c | $D_c = 0.3D_0$ | Patent pl. arch | Obstr. pl. arch |
| 1 – 6d | $D_d = 0.05D_0$ | Patent pl. arch | Obstr. pl. arch |

## 3. Results

We report in Table 4 the baseline perfusion values in each foot angiosome for the lesion-free scenario (Case 0) and their percent variations computed for all considered pathological scenarios (Cases 1-6) with respect to Case 0. Table 5 reports the time to peak (TTP) after cuff release in each foot angiosome for Case 0, and its percent variations computed for all considered pathological scenarios compared to Case 0, since longer TTPs can be associated with CLTI. Percent variations are computed as

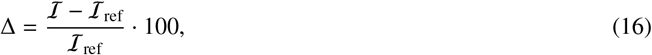

where ℐ is the considered quantity, and ℐ_ref_ is the corresponding reference value obtained for Case 0. Figures 3 and 4 show how cardiac-cycle averaged perfusion values vary in time in the DPA, LPA and MPA angiosomes for all considered scenarios.

**Table 4:**
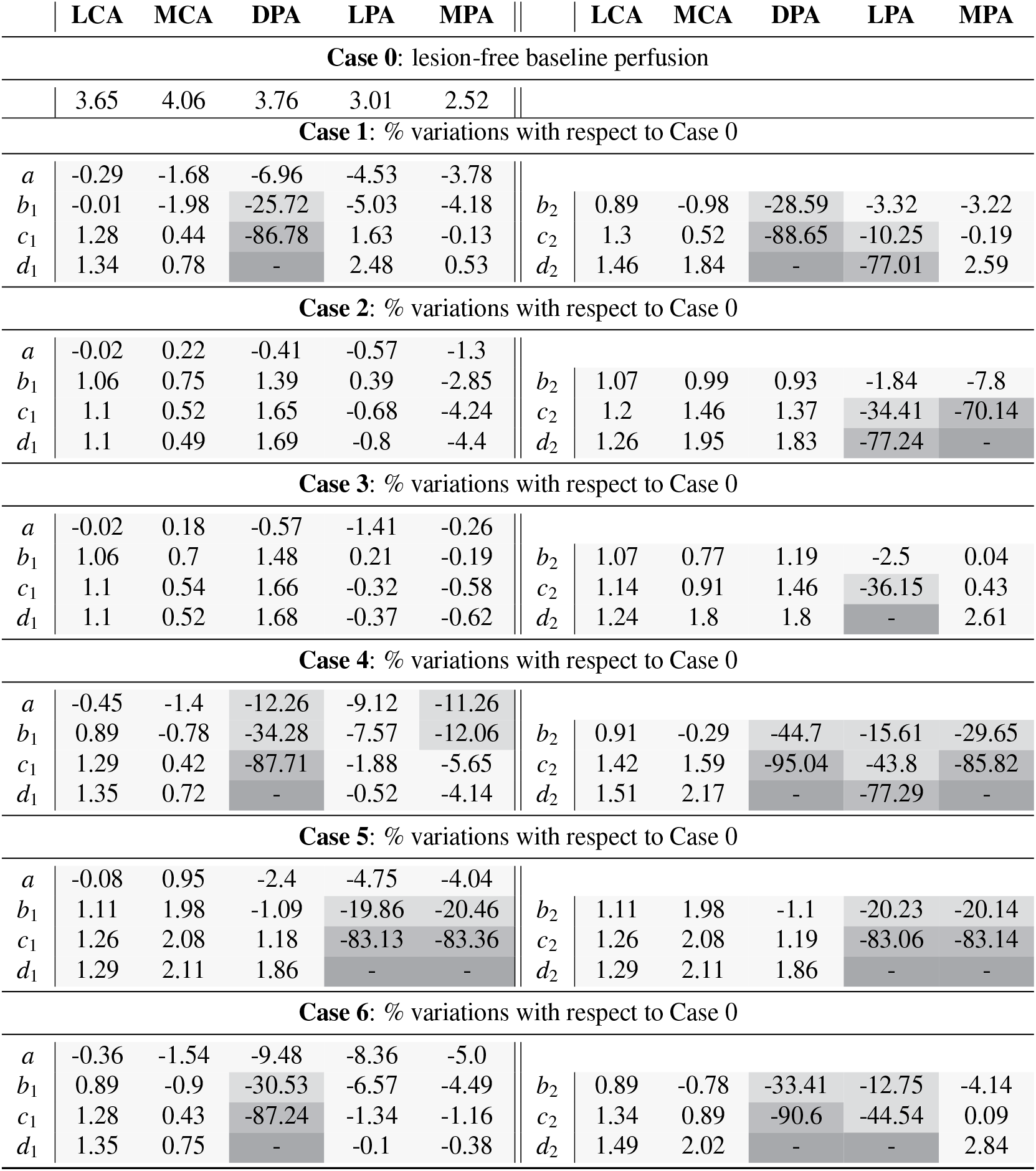
Baseline perfusion values in the foot angiosomes for a lesion-free baseline scenario (Case 0) and percent changes in baseline perfusion values compared to a model with no occlusion or stenosis (Cases 1-6). Cases 1*a*− 6*a* denote scenarios where all collaterals are assumed to be patent. Cases 1*b*_1_− 6*b*_1_, 1*c*_1_ −6*c*_1_, 1*d*_1_ −6*d*_1_ denote cases where the plantar arch is assumed to be fully patent, while other collaterals have a diameter that is, respectively, 60%, 30%, and 5% of the original diameter. Cases 1*b*_2_ −6*b*_2_, 1*c*_2_ −6*c*_2_, 1*d*_2_ −6*d*_2_ denote cases where all collaterals (including the plantar arch) have a diameter that is, respectively, 60%, 30%, and 5% of the original diameter.

**Table 5:**
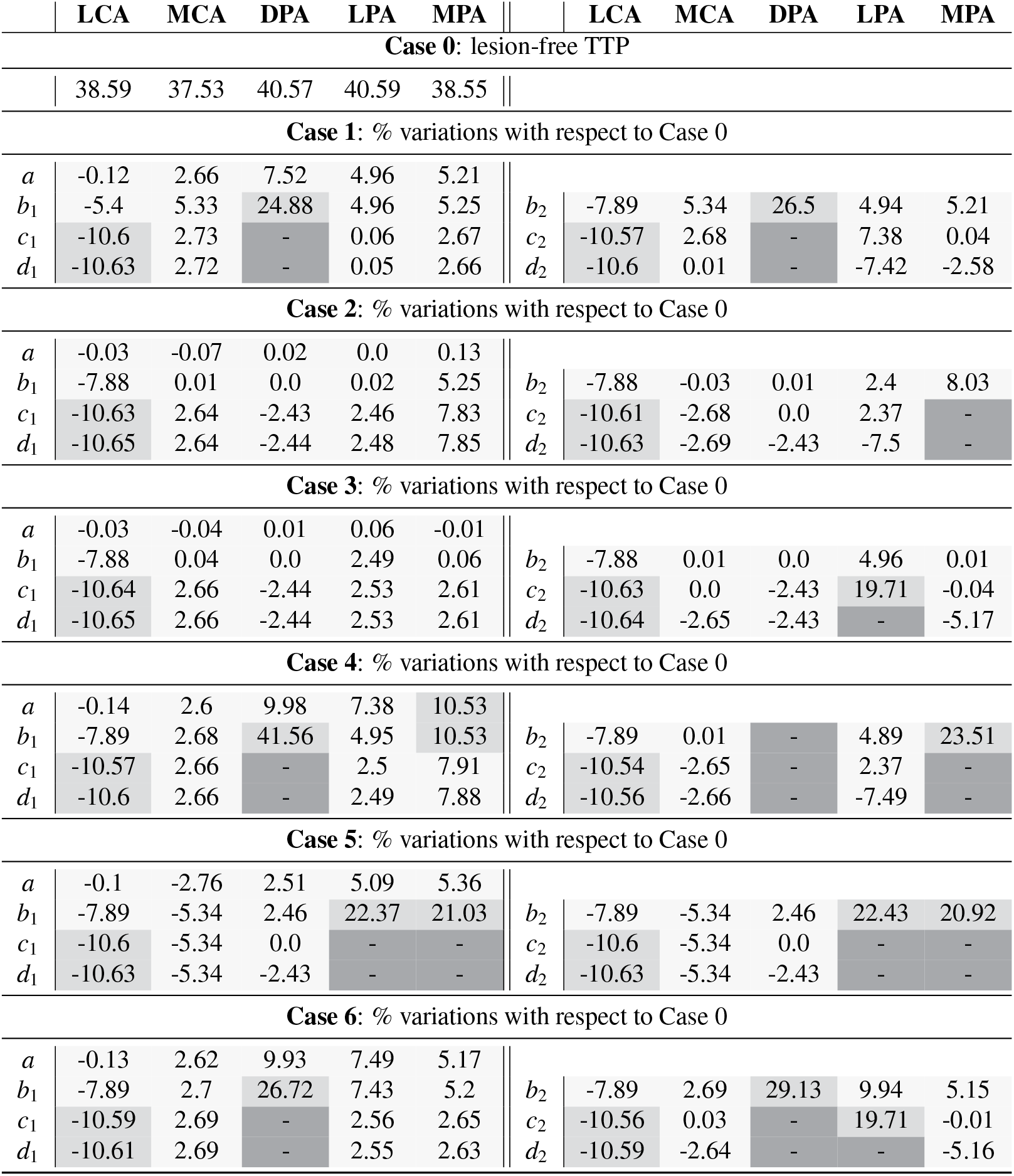
Time to peak after cuff release in the foot angiosomes for a lesion-free baseline scenario (Case 0) and percent changes in time to peak compared to a model with no occlusion or stenosis (Cases 1-6). If the peak did not occur within the first 75*s* after cuff opening, we assume that no peak occurs. Cases 1*a* −6*a* denote scenarios where all collaterals are assumed to be patent. Cases 1*b*_1_ − 6*b*_1_, 1*c*_1_ − 6*c*_1_, 1*d*_1_ − 6*d*_1_ denote cases where the plantar arch is assumed to be fully patent, while other collaterals have a diameter that is, respectively, 60%, 30%, and 5% of the original diameter. Cases 1*b*_2_ − 6*b*_2_, 1*c*_2_ − 6*c*_2_, 1*d*_2_ − 6*d*_2_ denote cases where all collaterals (including the plantar arch) have a diameter that is, respectively, 60%, 30%, and 5% of the original diameter.

**Figure 3.**
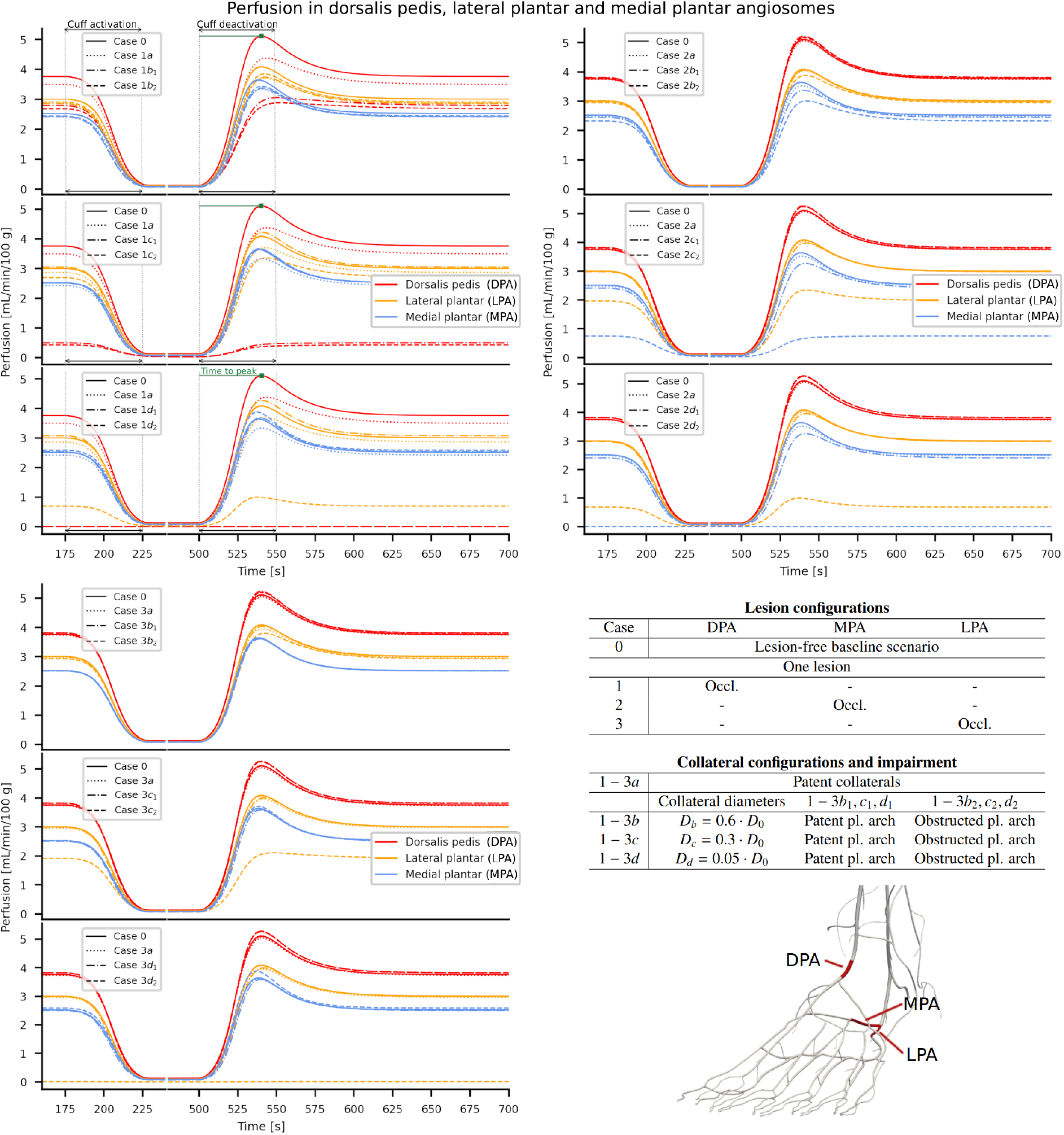
Time variation of cardiac-cycle averaged perfusion values in the DPA, LPA and MPA angiosomes for cases 1-3. Cases 1*a*− 3*a* denote scenarios where all collaterals are assumed to be patent. Cases 1*b*_1_ − 3*b*_1_, 1*c*_1_ − 3*c*_1_, 1*d*_1_ − 3*d*_1_ denote cases where the plantar arch is assumed to be fully patent, while other collaterals have a diameter that is, respectively, 60%, 30%, and 5% of the original diameter. Cases 1*b*_2_ − 3*b*_2_, 1*c*_2_ − 3*c*_2_, 1*d*_2_ − 3*d*_2_ denote cases where all collaterals (including the plantar arch) have a diameter that is, respectively, 60%, 30%, and 5% of the original diameter. In the top left panel, we highlight cuff activation, deactivation and the time to peak.

**Figure 4.**
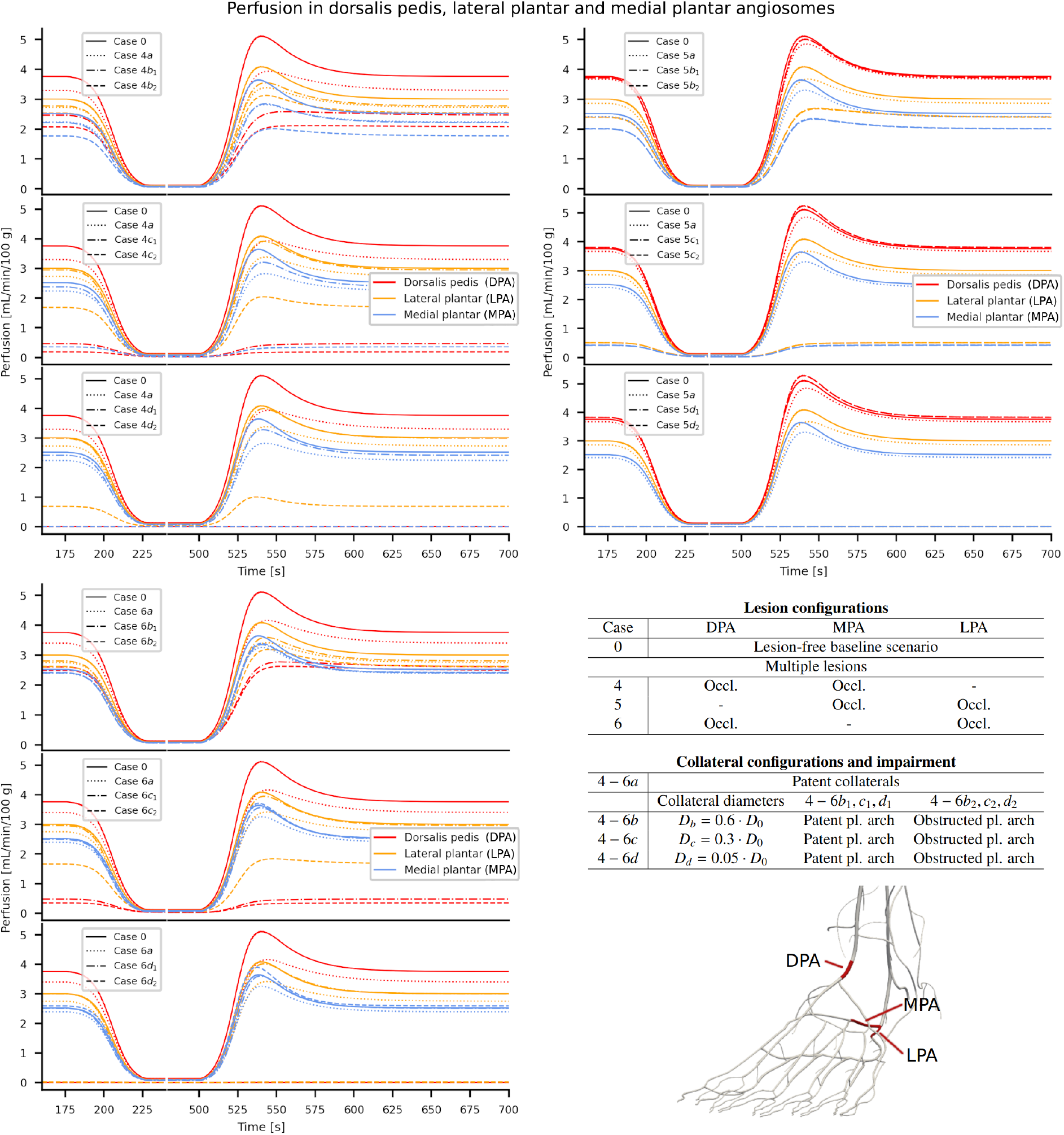
Time variation of cardiac-cycle averaged perfusion values in the DPA, LPA and MPA angiosomes for cases 4-6. Cases 4*a* − 6*a* denote scenarios where all collaterals are assumed to be patent. Cases 4*b*_1_ − 6*b*_1_, 4*c*_1_ − 6*c*_1_, 4*d*_1_ − 6*d*_1_ denote cases where the plantar arch is assumed to be fully patent, while other collaterals have a diameter that is, respectively, 60%, 30%, and 5% of the original diameter. Cases 4*b*_2_ −6*b*_2_, 4*c*_2_ 6−*c*_2_, 4*d*_2_ − 6*d*_2_ denote cases where all collaterals (including the plantar arch) have a diameter that is, respectively, 60%, 30%, and 5% of the original diameter.

## 4. Discussion

We discuss here the computational results presented in Section 3, highlighting their clinical implications to explore the capability of our model to characterise both the angiosome and “best-vessel” theories for revascularization.

### 4.1. Clinical interpretation and implications

The discussion will be focused on the impact of lesions in the DPA, LPA and MPA on their respective angiosomes. In Tables 4 and 5 we also report results for the LCA and MCA angiosomes, which are characterised by a low variation in baseline perfusion for all considered scenarios, due to the absence of lesions in their feeding arteries.

#### Perfusion under single-artery occlusion

Simulation results suggest that TTP in the angiosomes can serve as a functional readout of collateral sufficiency. Indeed, when collateralisation is preserved (collateral diameter ≥ 60% of the baseline geometry in cases 2 and 3, and not impaired in case 1), ΔTTP remains ≤10% in the affected angiosome under the condition of a single feeding artery occlusion. This pattern is consistent with maintained tissue perfusion through collaterals (Δ baseline perfusion ≤ 10%). In contrast, when the collateral calibre falls below a subcritical range (in case 1, 60% of the baseline diameter), the hyperaemic TTP response disappears, indicating that the autoregulatory capacity of the model was exhausted independently of cuff application due to the severity of the pathological condition. From a clinical perspective, this is consistent with the observed loss of perfusion. In cases 2 and 3, assuming all other arteries remain patent, TTP changes are primarily observed when the plantar arch is occluded, underscoring the arch’s role as the cross-angiosome conduit that distributes flow across MPA and LPA.

Placed in a clinical context, these findings offer a physiological bridge between the angiosome targeted strategy and the “best-vessel” paradigm. Prior observational studies have reported improved healing when revascularisation is directed to the artery feeding the wounded angiosome (Kim et al., 2021). They also observed that when the wound-bearing angiosome received blood flow via angiographically visible collaterals, outcomes were generally comparable to those achieved with direct angiosome perfusion (Špillerová et al., 2017; Chuter et al., 2024; Berchiolli et al., 2023). Our simulations explain how both strategies can be correct, conditional on arch patency and collateral sufficiency: if the plantar arch is patent and collaterals are functionally adequate (ΔTTP ≤ 10%), indirect or “best-vessel” revascularisation may restore sufficient perfusion to the wound angiosome via blood redistribution. If the arch is occluded, collaterals are absent, or the collateral diameter is, in our model, below 60% of its original value, simulations suggest that a direct angiosome-targeted revascularisation would be preferable. However, simulations are restricted to macrovascular architecture and vessel calibre, without incorporating microvascular dysfunction. In clinical reality, microvascular impairment, particularly in patients with diabetes, edema, or renal failure (Kim et al., 2021; Norgren et al., 2007), can significantly limit vasodilatory capacity and compromise collateral flow. This discrepancy between macrovascular and microvascular behaviour likely contributes to the observed variability in wound healing among patients with comparable angiographic findings.

#### Perfusion under dual-artery occlusion

When two of the three feeding arteries to the foot are occluded (cases 4-6), model predictions show a marked deterioration in perfusion, reflected by a substantial increase in TTP within the angiosomes with occluded inflow vessels, even when collateral diameters remain at 60% of the reference model for a healthy individual. When collateral diameter is reduced further to 30% or 5% of its original value, the TTP response is almost completely eliminated, indicating near-complete loss of functional perfusion, which is also confirmed by a reduction in baseline perfusion ≥ 40% in the affected angiosomes. This contrasts sharply with single-occlusion scenarios, where ΔTTP remained below 10% under collateralisation with a diameter above 60% of the standard model, suggesting that the physiological reserve provided by collaterals is insufficient once inflow is severely restricted. Interestingly, whether the plantar arch is patent or occluded only has a minimal effect in these dual-occlusion scenarios, underscoring that the dominant determinant of perfusion failure is the severe reduction in inflow rather than redistribution capacity. From a clinical perspective, in such scenarios, reliance on indirect strategies, even in the presence of visible collaterals, may be insufficient to achieve wound healing or limb salvage.

### 4.2. Limitations

The present study is characterised by several limitations. In particular, we perform a purely computational study employing an idealised vascular configuration, derived from clinical textbooks, which was modified to reproduce the application of a pressure cuff and a set of idealised pathological scenarios. These scenarios do not account for the various stenosis degrees that could be observed in clinical practice, nor for microvascular impairment, which is a fundamental flow-limiting factor, especially in the presence of comorbidities such as diabetes and oedema. Moreover, we reproduce the expected hyperaemic response after cuff release through a very simplistic description of control mechanisms. For these reasons, our model is currently not suitable for providing patient-level outcomes, and the findings reported in this study may not directly translate to patients with multilevel disease, severe infection/oedema, or extensive tissue loss.

## 5. Conclusions

In this work, we presented a computational model that can satisfactorily capture the perfusion dynamics in the angiosomes of the foot following a cuff-induced ischemia test. In particular, we used this model to study the impact of 42 pathological scenarios, characterized by occlusions placed in either the DPA, the MPA, or the LPA, and by different degrees of collateral impairment. If collateralization between foot angiosomes is preserved, occlusions in one of the three feeding arteries of the foot have little influence on overall foot perfusion, underlining the importance of blood redistribution in maintaining adequate perfusion levels. In this case, good patient outcomes can be achieved by revascularising the best-preserved vessel that crosses the ankle. When collateral vessels are impaired, instead, perfusion decreases in the angiosome corresponding to the occluded feeding artery. Additionally, perfusion in the MPA and LPA angiosomes is highly dependent on the level of patency of the plantar arch, which acts as a cross-angiosome conduit. In this case, it is important for the surgeon to revascularise the relevant feeding artery. Instead, when two of the three feeding arteries to the foot are occluded, there is a significant reduction in perfusion even at low collateral impairment levels, caused by the insufficient inflow. These considerations highlight the strong influence of network structure on simulation results, suggesting that meaningful comparisons with clinical data will require an accurate definition of the patients’ foot vasculature or, at least, a characterization of the degree of collateral impairment.

In the future, we plan to incorporate an explicit description of foot oxygenation in our model, as well as a characterisation of the variability in vascular properties caused by comorbidities commonly associated with peripheral artery disease. Moreover, we plan to study the impact of bypass surgeries on foot perfusion.

## Acknowledgments

C.D., and L.O.M. are members of the “Gruppo Nazionale per il Calcolo Scientifico” dell’ Istituto Nazionale di Alta Matematica (INdAM-GNCS, Italy). P.J.B. is supported by Brazilian agencies: CNPq (304347/2023-0) and FAPERJ (grant number E-26/200.364/2023).

## Data and code availability

The data and codes that support the findings of this study are available on request from the corresponding author. The complete model dataset is publicly available at http://hemolab.lncc.br/adan-web. The geometry and topology of the arterial network is described by the first 4041 lines (first vessel *A*_*celiac*_*trunk*_*C*_0, last vessel *P*_*intramedullary*_*T* _9_*ORG*29_*T* 9_*L*_4040) of the Supplementary file Data Sheet 1.CSV available at https://www.frontiersin.org/articles/10.3389/fphys.2023.1162391/full#supplementary-material. Model parametrization is outlined in (Blanco et al., 2014a; Müller et al., 2023).

